# FLUTE: a Python GUI for interactive phasor analysis of FLIM data

**DOI:** 10.1101/2023.03.31.534529

**Authors:** Dale Gottlieb, Bahar Asadipour, Thi Phuong Lien Ung, Chiara Stringari

## Abstract

Fluorescence lifetime imaging microscopy (FLIM) is a powerful technique used to probe the local environment of fluorophores. The phasor approach to FLIM data is a fit-free analysis and is increasingly used due to its ease of interpretation. To date, no open-source graphical user interface (GUI) for phasor analysis of FLIM data is available thus limiting the widespread use of phasor analysis in biomedical research. Here we present (F)luorescence (L)ifetime (U)l(t)imate (E)xplorer (FLUTE), a Python GUI that is designed to fill this gap. FLUTE simplifies and automates many aspects of FLIM analysis, such as calibrating the FLIM data, performing interactive exploration of the phasor plot with cursors, displaying the phasor plot and the FLIM images with different lifetime contrasts and calculating the relative concentration of molecular species. The final edited datasets after applying the desired filters and thresholds can be exported for further user specific analysis. FLUTE was tested using several FLIM datasets including autofluorescence of Zebrafish embryos, cells in vitro and intact live tissues. In summary, our user-friendly GUI extends the advantages of phasor plotting by making the data visualization and analysis easy and interactive, allows for analysis of large FLIM datasets and accelerates FLIM analysis for non-specialized labs.

**Impact statement:** This work introduces the first open-source graphical user interface (GUI) for phasor analysis of Fluorescence Lifetime Microscopy (FLIM) data. Phasor analysis is increasingly used for FLIM data analysis in biomedical research as it reduces the complexity of the analysis and provides a powerful visualization of the data content and optimization of data handling with respect to multiexponential fitting. However, the development of quantitative FLIM applications in the life sciences has been until now hampered by the lack of an open source and user-friendly graphical user interface. Here we introduce FLUTE that expands some possibilities of phasor FLIM image processing, accelerates the whole FLIM analysis and simplifies the visualization and the analysis of FLIM data, thus making phasor analysis possible for a broader base of researchers. FLUTE will be of interest to researchers with interests ranging from physics to biology and will facilitate research in several biomedical fields.

## 1. Introduction

Fluorescence lifetime imaging microscopy (FLIM) is a powerful technique used to probe the local environment of fluorophores reporting on pH, fluorescence resonance energy transfer *(FRET)*, binding and metabolism [1–6]. FLIM is increasingly used in biology and biophysics applications because the lifetime of a fluorophore can be used to provide functional contrast in images. However, the use of quantitative, easy, fast and interactive analysis of large FLIM datasets has been until now hampered by the lack of suitable open source software.

The multi fluorescence decay curve that is measured in every pixel of the image can be analyzed by fitting the decay with multi-exponential function [3, 5]. Several companies have developed commercial closed source packages for multi-exponential FLIM analysis that usually support their own file format [7] and recently some free and open source software has been developed for exponential fitting such as FLIMfit [8], FLIM –FRET analyzer [9], Flimview [10] and FLIMJ [11].

An alternative to multi exponential curve fitting that has been gaining popularity is the phasor analysis to FLIM [12–14] as it reduces the complexity of the analysis while at the same time providing a powerful visualization of the data content and optimization of data handling [14–16]. Phasor analysis is a fit-free technique in which the fluorescence decay from each pixel is transformed with a fast Fourier transform (FFT) into a point in two-dimensional phasor space. The main advantages of phasor analysis are that it provides a visual distribution of the molecular species by clustering pixels with similar decays, it allows for colour mapping of the pixels of the FLIM images, and it is linear in terms of non-interacting molecular species [12, 13, 17, 18]. Phasor analysis of FLIM data is increasingly used in several fields of scientific research [19–24] and is particularly used for metabolic imaging with the endogenous biomarkers NAD(P)H and FAD to map their complex autofluorescence distributions in live tissues [5, 13, 25–33].

The most commonly used software in biomedical research for phasor analysis is SimFCS which has been developed by the Laboratory for Fluorescence and Dynamics [14]. While companies have been increasingly developing phasor interfaces to analyze their FLIM data acquired with their systems, to date, an open-source and easy to use software for phasor analysis of FLIM data is not available.

Here, we developed (F)luorescence (L)ifetime (U)l(t)imate (E)xplorer (FLUTE) that is designed to fill this gap, and allow simple, quick, and interactive visualization of FLIM data as well as advanced analysis using phasors. The user-friendly graphical user interface (GUI) expands some limited possibilities of phasor FLIM image processing, accelerates the FLIM analysis pipeline and allows for analysis of large FLIM datasets. FLUTE can be used by microscopy labs to perform real time phasor analysis during their experiment and also allows for post processing by exporting the data. It is designed to be user-friendly for researchers of different backgrounds who may not be specialized in FLIM.

## 2. Results

### 2.1 GUI Development and Implementation

FLUTE is programmed in Python using PyQt5 [34] for the graphical user interface (GUI) while and phasor analysis (Methods 3.1) and date processing is performed using NumPy [35] and SciPy libraries [36] (Fig1). Both the Python code and the executable file are free and open-source (See supplementary material and Fig S1 and S2). The easiest way to use FLUTE is to run the available ‘FLUTE.exe’ file, which has been tested to work on Windows 7 and 10, and doesn’t require Python to be installed. ‘main.py’ can also be run after installing Python (tested with Python 3.10). FLIM data is read as a tiff stack format (Fig2), where each image of the stack represents a temporal bin of the FLIM stack acquired in the time domain.

**Fig. 1.**
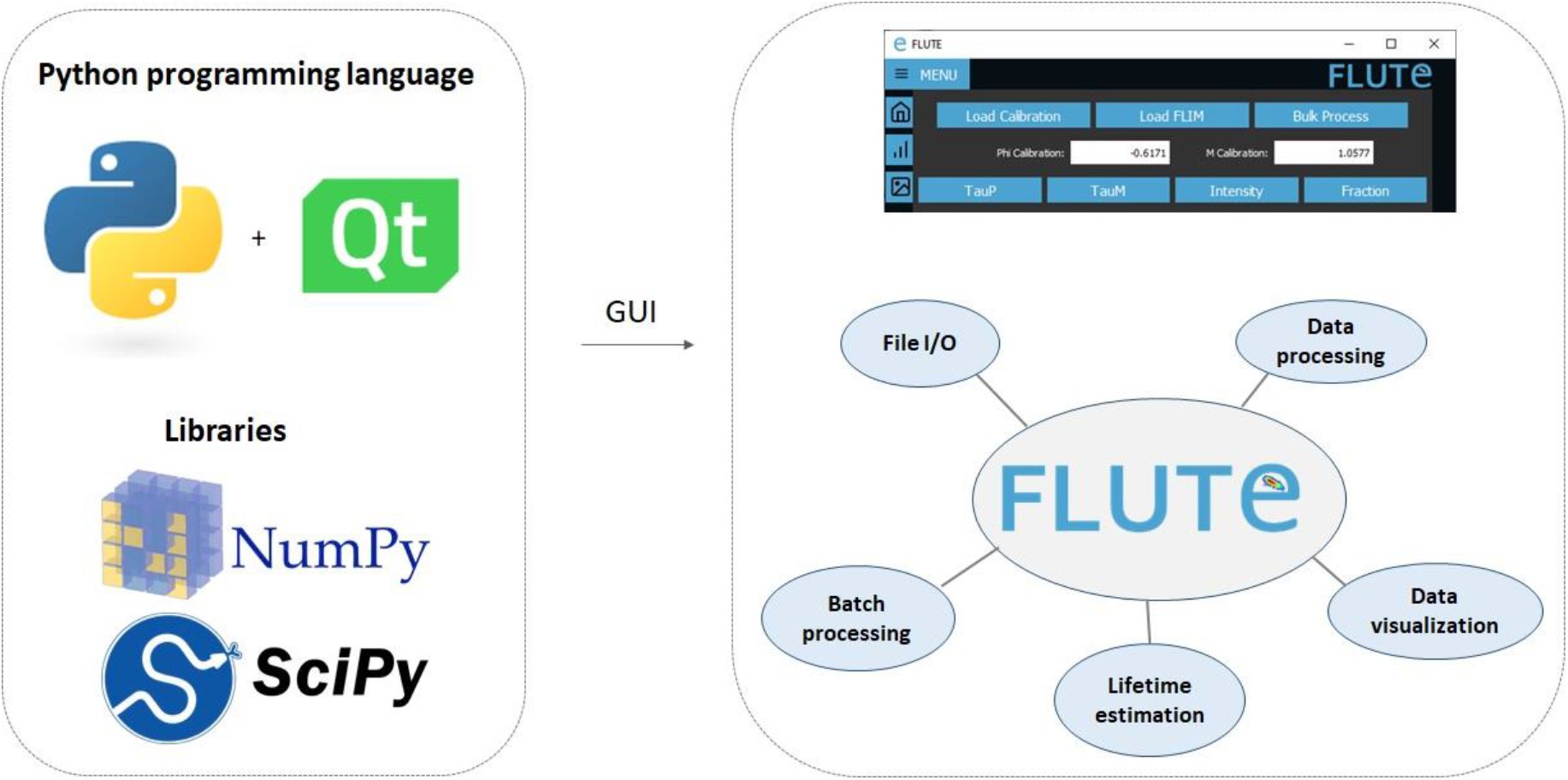
FLUTE language and architecture. FLUTE is programmed in Python using PyQt5 and using NumPy and SciPy libraries. The Graphical user Interface is designed to be user-friendly and to perform different functionalities such as import and save data, data processing, data visualization, lifetime estimation and batch processing.

**Fig. 2.**
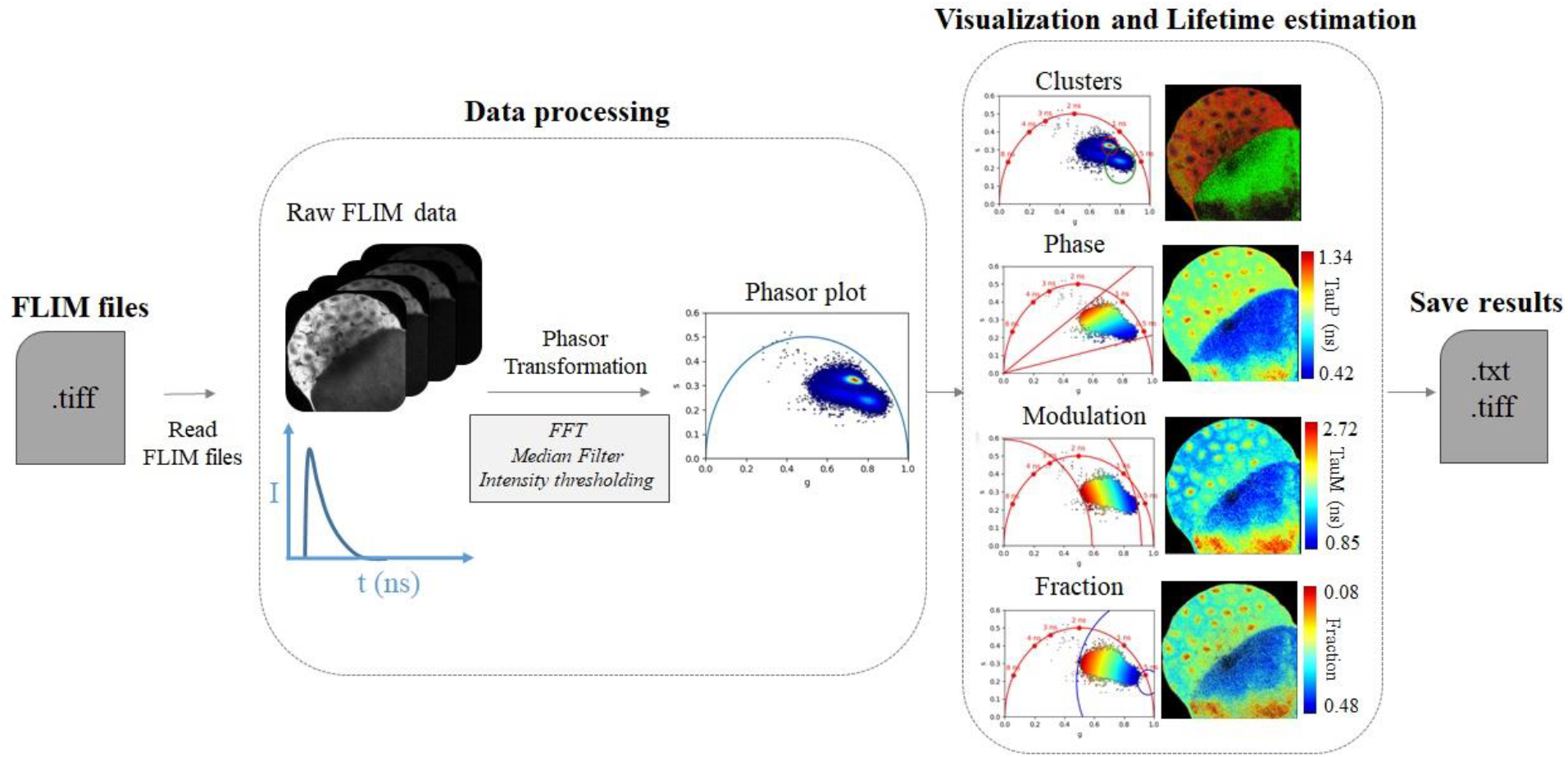
FLUTE processes and tasks. The architecture of FLUTE has been designed and optimized to perform user-friendly and efficient Data processing and data visualization and exploration as well as to easily open FLIM data and save results.

The FLUTE architecture has been designed to perform several tasks and processes in a user-friendly way (Fig1):

#### File I/O

- Read FLIM files (.tiff)
- Saves results to files (analysis and figures)

#### Data processing

- Calibration (with a reference standard)
- Phasor transformation at different harmonics
- Filters (median filter, thresholds,..)
- Thresholding (ROI identification)

#### Data Visualization

- Interactive Exploration of data with cluster analysis
- Displaying the phasor plot and FLIM images with different lifetime contrast

#### Lifetime estimation

- Calculation of tauPhase and tauModulation lifetimes
- Calculation of fraction of molecular species (i.e. bound NADH)
- Calculation of average lifetime in ROIs

#### Batch processing

- Processing of large FLIM datasets

### 2.2 FLIM Data Processing

FLUTE has been programmed to perform the phasor analysis of the FLIM data (Methods 3.1) acquired in the time domain through an FFT (Fig. 2). The phasor coordinates g and s are calculated by using equations 1 and 2 (Methods 3.1). The main window of FLUTE is shown in Fig. 1 and Fig S3. FLIM data is converted to the phasor plot with an FFT and referenced using a fluorescence calibration standard of known single lifetime (Fig. 2 and Fig. S4), such as Fluorescein with lifetime 4ns [14] (See Methods and Fig. 5). FLUTE allows analysis of FLIM data acquired with different parameters such as temporal bin number, bin width and laser repetition rates that need to be specified in the GUI (Fig. 2 and Fig. S4).

**Fig. 3.**
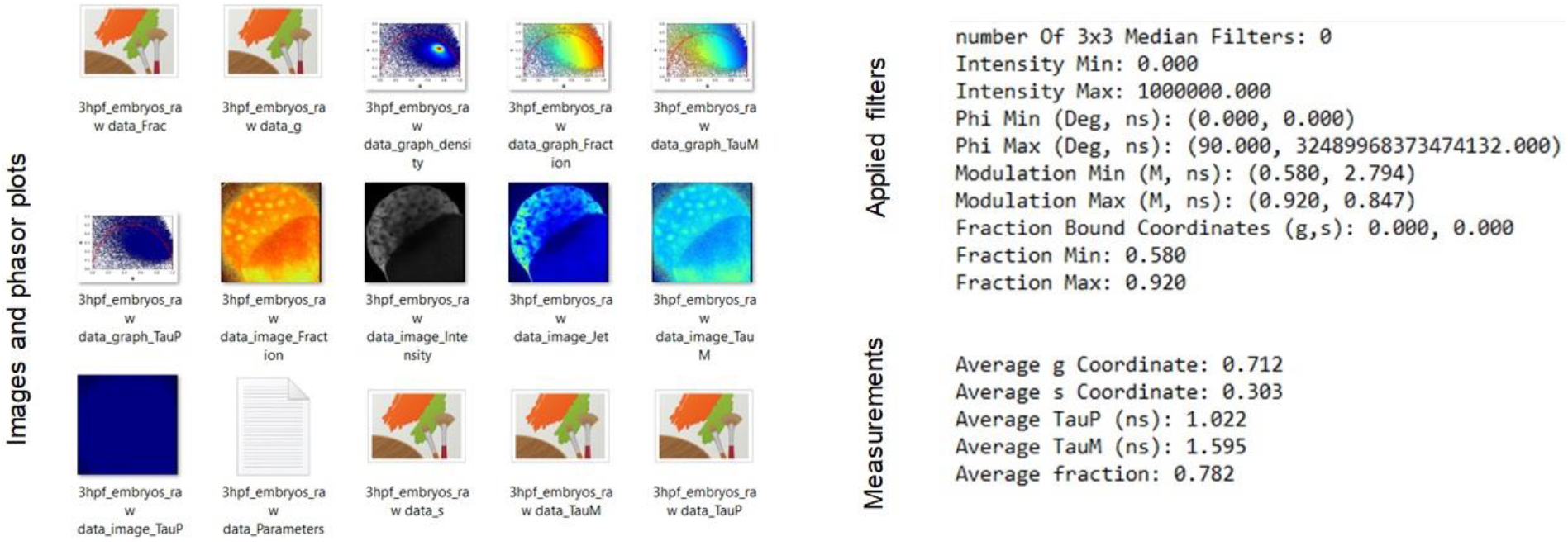
FLUTE saved results. Saved FLIM images and phasor plots (left) and applied filters to create the mask and measurements of the average of g, s, TauPhase (TauP), TauModulation (TauM) and fraction (right).

**Fig. 4.**
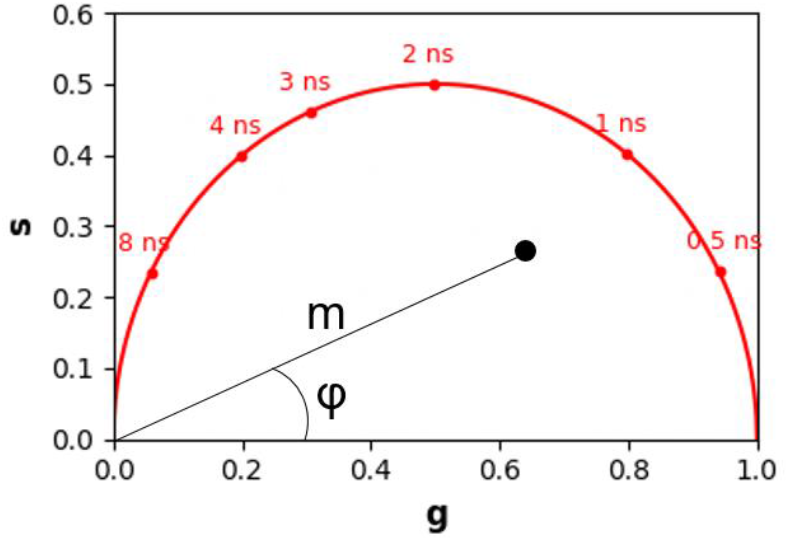
Phasor plot representation. Multi-exponential decay is located within the universal circle of the phasor plot. The phasor coordinates g and s are the real and imaginary part of the FFT.

**Fig. 5.**
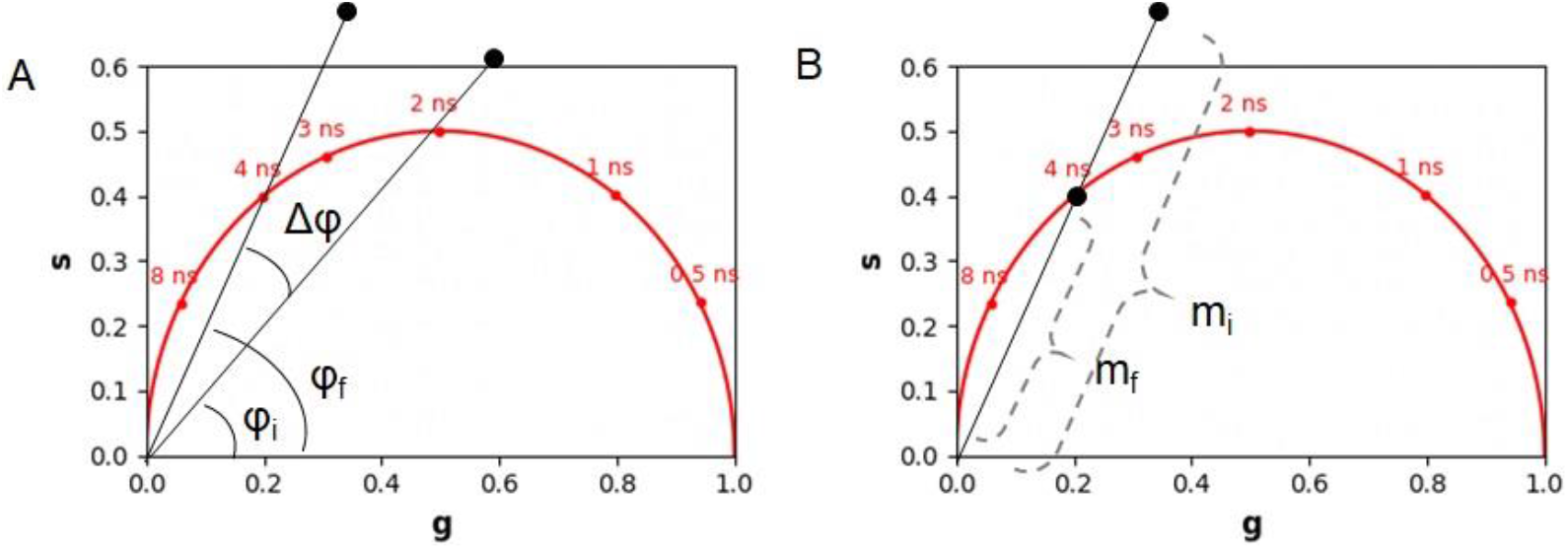
Calibration of the phasor plot using a fluorescent standard of fluorescein of 4 ns. (A) Phase correction (B) modulation correction.

When FLIM data is first opened (Fig. S5), the initial phasor plot is displayed and colour mapped as a histogram showing the density of points in phasor location and the initial intensity image is calculated as the sum of the .tiff stack (Fig. S5). FLUTE allows to apply a 3×3 convolutional median filter on the phasor g and s coordinates (Fig. S7) to reduce the spread of phasor points and reduce the noise without affecting the resolution of the image [17] and an intensity filter through a threshold (Fig S8).

### 2.3 FLIM data visualization and lifetime estimation

We designed FLUTE to be a user-friendly GUI for flexible data exploration and easy data visualization (Fig. 2). Interactive exploration of the FLIM data can be performed by using multiple coloured cursors of variable sizes to select pixels with similar fluorescence decays and phasor locations, highlighting simultaneously the corresponding pixels in the image (Fig. 2, S16 and Supplementary Information 8.4).

FLUTE also allows to estimate the lifetime of the FLIM image [12] and to choose the appropriate colour mapping of phasor plot and FLIM images for data exploration (Fig S3 and S9). The threshold of these colour maps for TauPhase (TauP) and TauModulation (TauM) can be adjusted either via the Interactivity window (Fig. S10) or via the Thresholding window (Fig. S11) of the GUI.

Mapping of the FLIM image and of the phasor plot according to lifetime contrasts such as TauPhase or TauModulation is performed simultaneously allowing a clear representation and an interactive exploration of the FLIM data (Supplementary Information 8.1 and 8.2). TauPhase is estimated with equation 7 and TauModulation with equation 8, respectively.

If the FLIM image contains a combination of two fluorescent non-interacting molecular species (i.e. excluding FRET) that give rise to a linear combination in the phasor space (Fig. 6), FLUTE can calculate and map the faction of one molecular species by either inserting the single lifetime or their phasor coordinates of the second species (Fig. 6, Fig. S14 and Supplementary Information 8.3). For example, the fraction of bound/free NADH can be calculated and mapped graphically in every pixel from the location of the free NADH that has a known single lifetime of 0.4 ns [37] (Fig. 2 and S15) using equation 13.

**Fig. 6.**
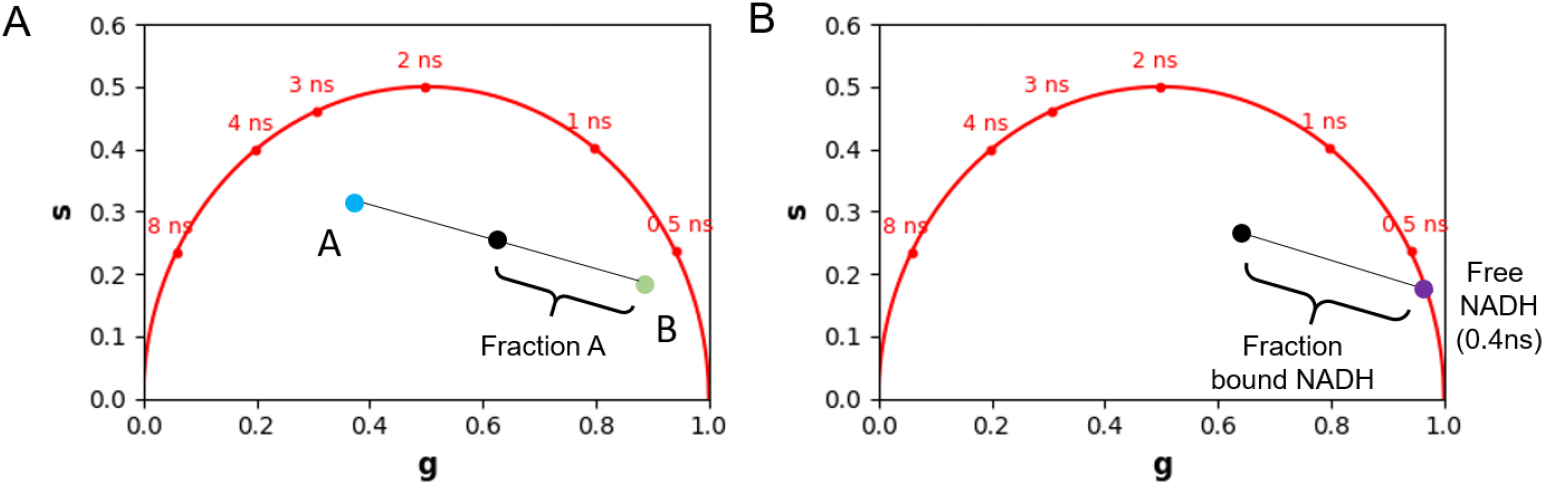
Graphical calculation of fraction of molecular species in a linear combination. (A) Calculation of fraction the molecular species A (B) graphical calculation of fraction of bound NADH.

### 2.3 Saving results and batch processing

We implemented image masking by applying thresholds on the lifetime values or on the fraction of the molecular species. Once the desired thresholding is applied on the FLIM image (Fig. S17), the data with the applied mask can be saved (Fig.2, Fig3 and Supplementary Information 9.1). FLUTE saves the matrices of phasor coordinates, the lifetime contrasts and the fraction of the molecular species in tiff files as well as the calculated corresponding average values in the mask (Fig.3).

Finally, FLUTE allows batch processing on multiple FLIM images analyzed using the same parameters (Fig. S20 and Supplementary Information 10).

## 3. Methods

### 3.1 Phasor analysis

The multi-exponential intensity decay of every pixel of the FLIM image acquired in the time domain is converted in the phasor plot with a Fourier transform using the following formulas:

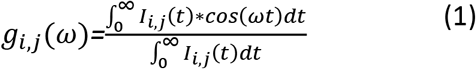

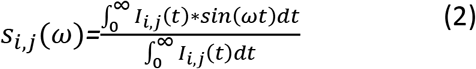

Where *i* and *j* indicate the order pixel of the image and ω is the frequency. ω is calculated through the laser repetition rate *ω*=*2πf*. The real (*g*) and imaginary (*s*) parts are plotted in the graphical phasor plot (Fig. 4). In the frequency domain, the FLIM data in the frequency domain can be expressed as follows:

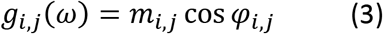

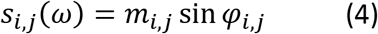

Where the phase *φ*_i,j_ and modulation *m_i_*, are represented in Fig. 4 and calculated using the following equations:

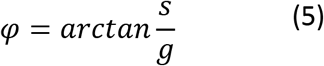

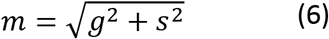

Estimations of the lifetime in each pixel in terms of the phase and modulation can be performed by:

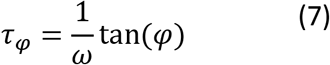

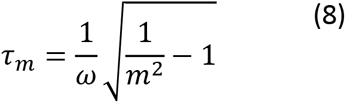

Phase (*φ*) and modulation (m) of the phasor cloud are first referenced using calibration to account for the instrument response function and the delays of the electronics. The calibration is performed with a reference sample that is usually a fluorophore with a known fluorescence lifetime, for example, fluorescein that has a decay of 4 ns or with an instrument response function like the Second-harmonic Generation measurement of 0ns. An example of fluorescein decay is uploaded in the supplementary file as *Fluorescein.tiff* file. Fig. 5 demonstrates the principle of the phasor calibration with Fluorescein: a correction on the phase is performed applying an offset of Δ*φ*, while correction on the modulation is performed applying a multiplication constant Δ*m* as described in the following formulas:

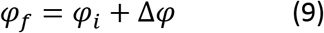

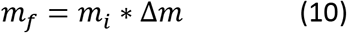

If the FLIM image contains a combination of two non-interacting fluorescent molecular species (i.e. excluding FRET) it gives rise to a linear combination in the phasor space (Fig. 6). In a system with two fluorescent species A and B the phasor the experimental point (black) lies along a straight line joining the phasors of the two species A and B.

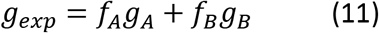

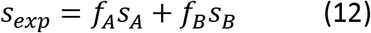

Where *f_A_* is the fractional contribution of molecular speaie A and *f_B_* is the fractional contribution of molecular species B.

The fraction of molecular species A (f_A_) is graphically calculated as the distance between the black experimental point *g_exp_*, *s*_exp_) and the location of the molecular species B (*g_B_*, *s*_B_) using the following equation:

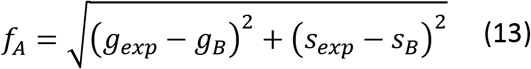

### 3.2 Two-photon excited fluorescence lifetime microscopy imaging of zebrafish embryo

Fluorescence lifetime microscopy imaging was performed with a laser scanning microscope (TriMScope, Lavision Biotec, Germany) equipped with a dual-output femtosecond laser (Insight DS++, Spectra-Physics, Santa Clara, CA, USA) with a tunable beam from 680 nm to 1300 nm (120 fs pulses, 80 MHz) and a fixed beam at 1040 nm (200fs pulses). The excitation laser is focused on the sample through a water immersion objective (25X, NA=1.05, XLPLN-MP, Olympus, Japan). The fluorescence signal is collected through the same objective and then epi-detected by a hybrid photomultiplier tube (R10467U, Hamamatsu, Japan). FLIM is performed with a Time-correlated single photon counting (TCSPC) electronics (Lavision Biotec, Germany) with 5.5 ns dead time, and 27 ps time bins. The laser trigger is taken from the fixed wavelength beam using a photodiode (PDA10CF-EC, Thorlab). We used a fluorescein solution at pH9 with a single exponential of 4.04 ns to calibrate the FLIM system. We performed FLIM imaging of zebrafish autofluorescence with 740 nm excitation wavelength with a typical power of 25 mW and collected the fluorescence signal through a band pass filter Semrock FF01–460/80. We typically collected 400-800 photons during an acquisition time of 80 seconds for a 512×512 pixels image with a pixel dwell time of 162 μs/pixel (18 images with 9.04 μs/pixel are accumulated). We imaged a zebrafish embryo at three hours post fertilisation (3hpf) embedded in agarose.

## 4. Conclusion

Here we have presented FLUTE, a graphical user interface which removes barriers in FLIM analysis making the phasor approach more accessible for applications in biology and the biomedical field. FLUTE is a very easy to use and highly interactive GUI for FLIM data analysis as well as open-source. FLUTE simplifies and automates many aspects of the phasor FLIM data processing and visualization (Fig 2) such as calibrating the FLIM data, displaying the phasor plot, calculating the relative concentration of molecular species, evaluating lifetime, displaying the phasor plot clouds and the FLIM images with different lifetime contrast, performing interactive reciprocity analysis with cursors, applying different types of filters and thresholding and calculating average lifetime values in masks. The final edited datasets after applying the desired filters and thresholds can be exported for further user specific analysis (Fig 3). FLUTE allows simultaneous analysis of multiple images and it has also been designed to perform batch processing of large FLIM datasets. Overall, FLUTE expands some possibilities of phasor FLIM image processing, accelerates the whole FLIM analysis pipeline and simplifies the visualization and the analysis of FLIM data, thus making phasor analysis possible for a broader base of researchers.

## Supporting information

Supplementary information - User Manual

## Acknowledgements

The authors thank Ana Maria Pena for feedback on the GUI and Emmanuel Beaurepaire for the choice of the GUI name. This paper has been submitted to bioRxiv preprint (BIORXIV/2023/534529)

## Author Contributions

DG designed and implemented the GUI. CS designed and supervised the research. DG and CS wrote the manuscript. DG, TPLU, BA and CS tested the GUI. All authors read and approved the manuscript.

## Data Availability Statement

FLUTE is free to use under the GPL3.0 license. Source code and executable are available on GitHub page: https://github.com/LaboratoryOpticsBiosciences/FLUTE

## Supplementary Materials

To view supplementary material for this article, please visit the journal website

