## Supplementary information - User Manual for "FLUTE: a Python GUI for interactive phasor analysis of FLIM data"

### Supplementary material - User Manual

### FLUTe

#### Fluorescence Lifetime UlTimate Explorer

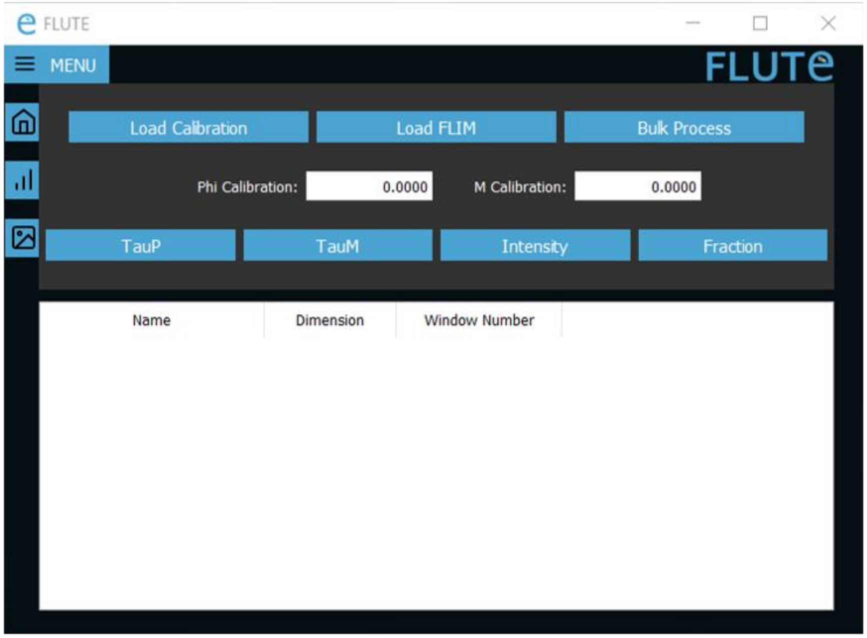


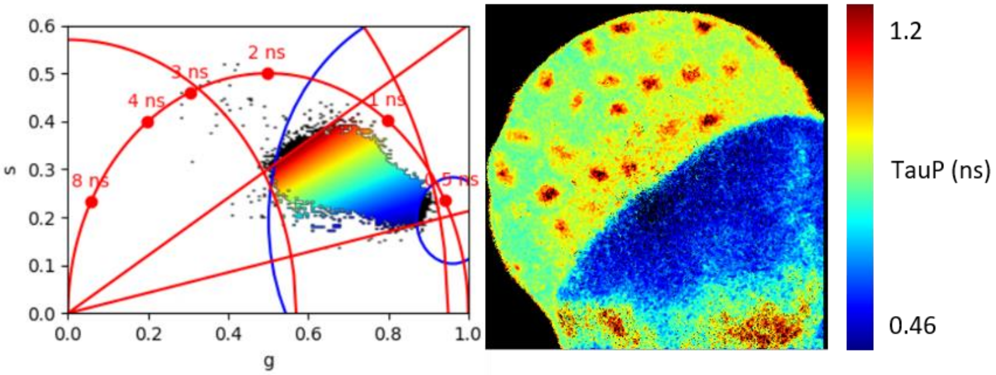

### Table of contents

1. Running FLUTE ... 3
   1.1 Run exe file ... 3
   1.2. Run main.py ... 3
2. Main window navigation ... 4
3. Calibration with a lifetime reference ... 4
4. Loading FLIM data ... 5
5. Selecting datasets using the table ... 5
6. Median filter ... 6
7. Intensity filter ... 6
8. Changing colour maps ... 7
   8.1 TauP colour map ... 8
   8.2 TauM colour map ... 8
   8.3 Fraction colour map ... 9
   8.4 Circle selection ... 10
9. Interactivity window ... 10
   9.1 Range adjustment ... 11
   9.2 Closing datasets ... 12
   9.3 Saving data ... 12
10. Batch processing ... 13

### 1. Running FLUTE

FLUTE can be run in two different ways.

#### 1.1 Run exe file

Download FLUTE.exe from GitHub (https://github.com/LaboratoryOpticsBiosciences/FLUTE) and double-click FLUTE.exe.

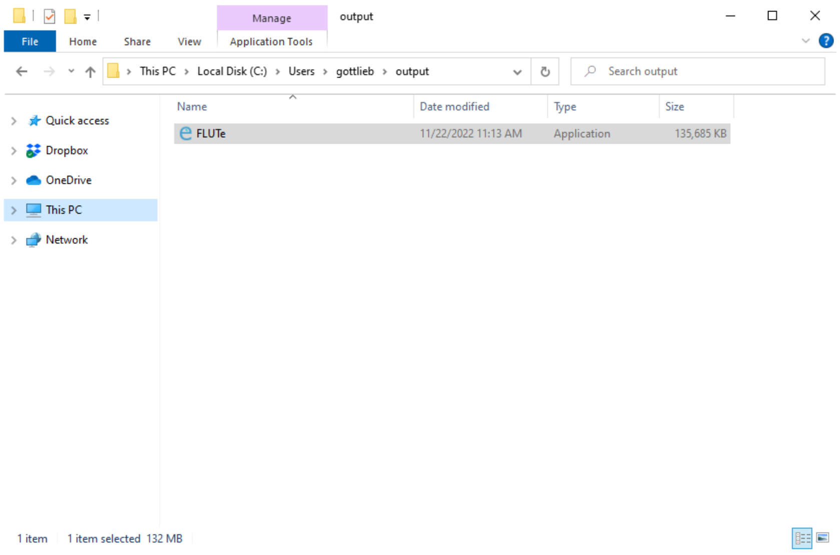


***Figure S1****: Running FLUTE.exe*

#### 1.2. Run main.py

If Python is installed on your system, open the terminal; navigate to the folder containing main.py, and type: python main.py. Install all required dependencies using pip.

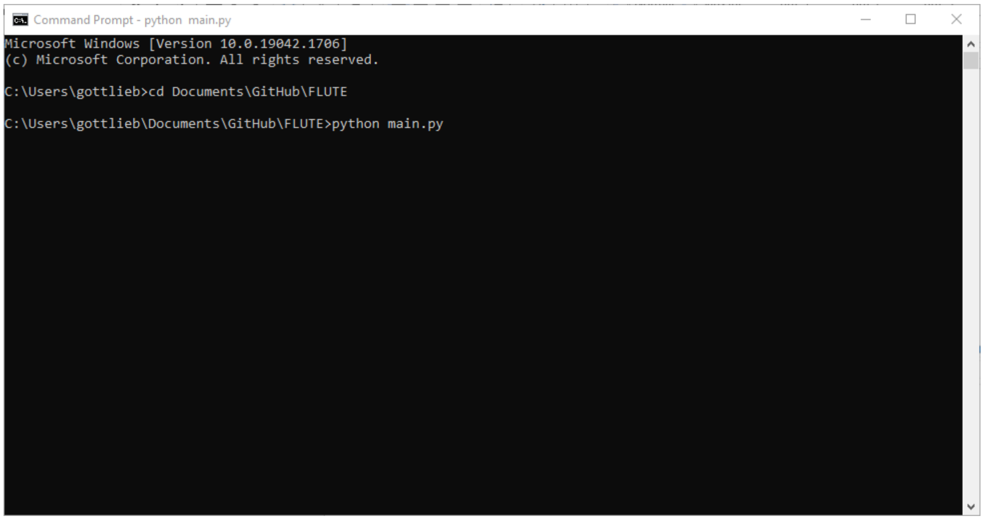


***Figure S2****: Running main.py using a computer with Python installed*

#### 2. Main window navigation

Figure S3 shows how the main window is divided into several sections, referred to as **Home**, **Interactivity**, **Thresholding**, and **Table** for the remainder of this document. The names of the windows are accessed by pressing the **Menu** button.

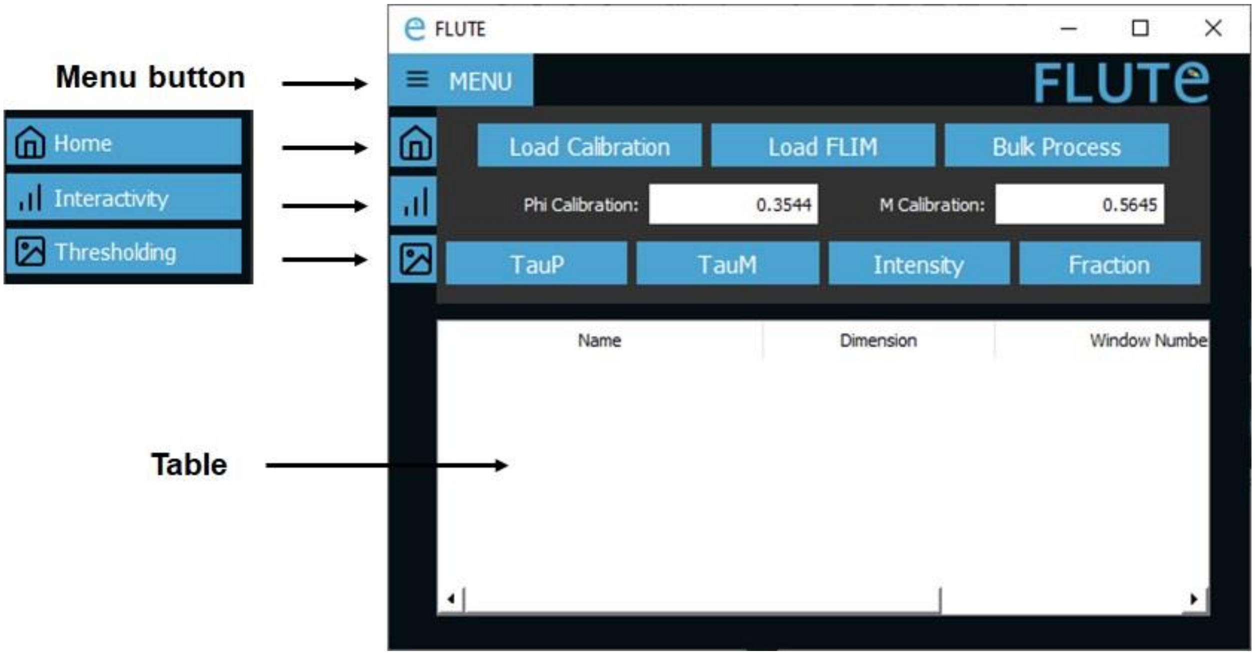


***Figure S3:*** *labeled windows for FLUTE main user interface. On the left, there are the Home window, Interactivity window, and Thresholding window buttons. Table displays the list of loaded data.*

#### 3. Calibration with Lifetime reference

To calibrate FLIM data with FLUTE, click **Load Calibration** on the **Home** window and enter the setup parameters in the popup window:

- Bin Width (ns): Duration of a single temporal bin of the time-domain FLIM acquisition
- Laser Freq. (MHz): Laser repetition rate
- Tau Ref. (ns): Known lifetime of the single-exponential reference sample (e. g. 4 ns for fluorescein)
- Harmonic: Integer multiple applied to laser frequency that will be used to calculate the Fourier Transform

FLUTE calculates the **Phi Calibration ($\Delta\varphi$)** and **M Calibration ($\Delta m$ )** values (equations 9 and 10 of main text) to be applied for calibration and displays them in the home menu as shown in Fig. 4. After calibration if you load the same calibration file (*Fluorescein.tiff)* file it is now located at the correct location of 4 ns in the phasor plot. The same **Phi Calibration ($\Delta\varphi$)** and **M Calibration ($\Delta m$ )** values will be applied to all the FLIM data opened with the **Load FLIM** function.

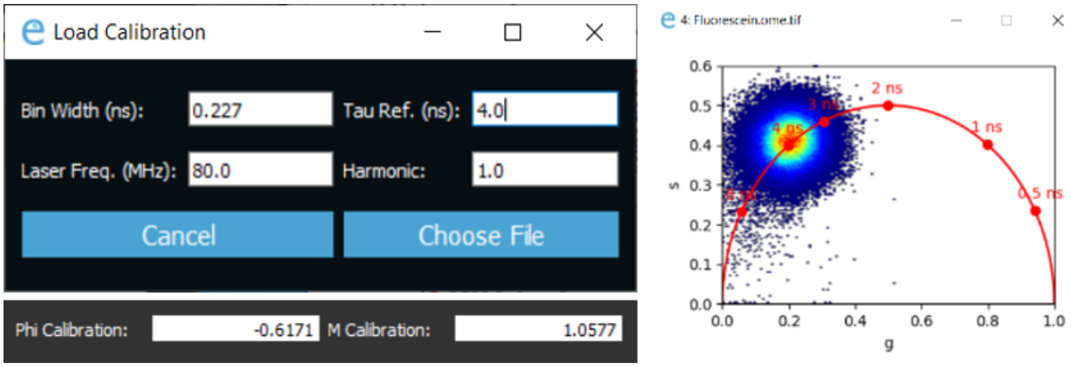


***Figure S4:*** *Calibration of FLUTE and example of calibrated experimental fluorescein sample with 4 ns lifetime.*

### 4. Loading FLIM data

In the **Home** window, press **Load FLIM**, and select the data to open. FLIM data must be uploaded as a .tiff stack. You can use the shortcut ctrl+click or shift+click to select multiple datasets at once. The phasor plot with a histogram density colour map and intensity image will open for each dataset, and the table will be populated. Figure S5 shows the example of *3hpf_embryos_raw data.tiff* that can be found in Supplementary data.

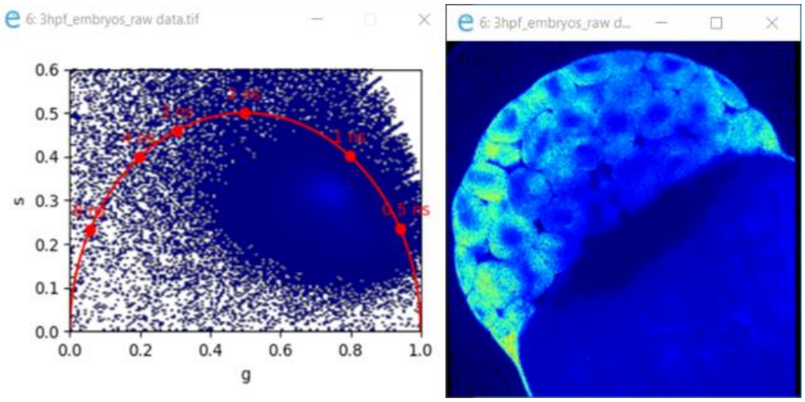


***Figure S5:*** *Loaded data 3hpf_embryos_raw data before any changes are applied.*

### 5. Selecting active datasets using the table

In all following sections, changes to datasets are only applied when the data is active and selected in the table. To select data, either click whichever is desired, or ctrl+click or shift+click to select multiple sets of data.

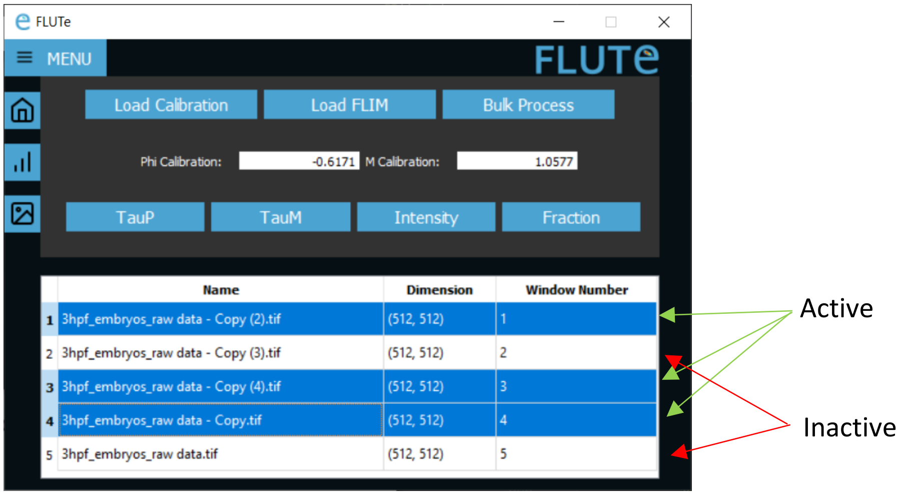


***Figure S6**: Selecting active and inactive data in the Table on the Main Window.*

In the example provided in figure S6, changing parameters such as colour maps or intensity threshold will only apply to windows 1, 3, and 4, but will not be applied to windows 2 and 5.

### 6. Median filter

Successive median filters can be applied to the g and s matrices using the **Median Filt** entry box within the **Thresholding** window. A 3x3 convolutional median filter is applied n times to the s and s coordinates of the phasor plot using scipy.signal.medfilt. Figure S7 displays the use of 3 median filters applied to the *3hpf_embryos_raw data.tiff* FLIM file three hours post fertilization (as for now referred to as 3hpf) data:

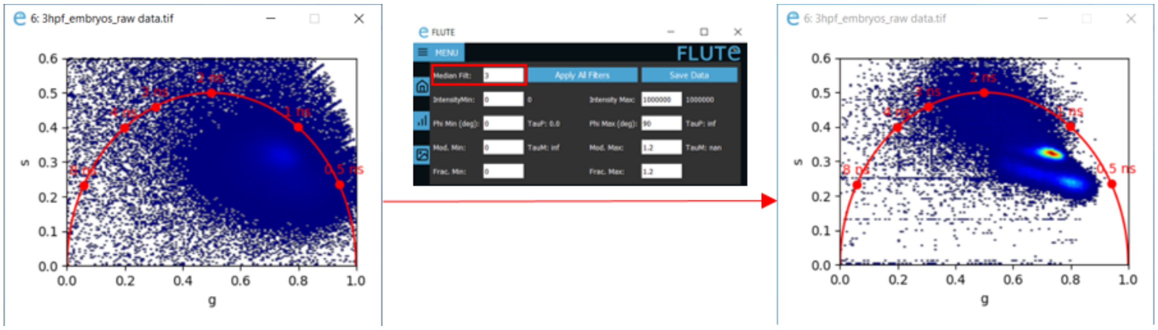


***Figure S7:** Applying 3x3 convolutional median filter to data.*

### 7. Intensity filter

Minimum and maximum intensity thresholds can be applied using the **IntensityMin** and **IntensityMax** entry boxes inside the **Thresholding** window. The intensity is calculated as the sum of all images in the .tiff stack. As shown in figure S8, applying a min intensity threshold of 25 counts to the 3hpf dataset removes the background:

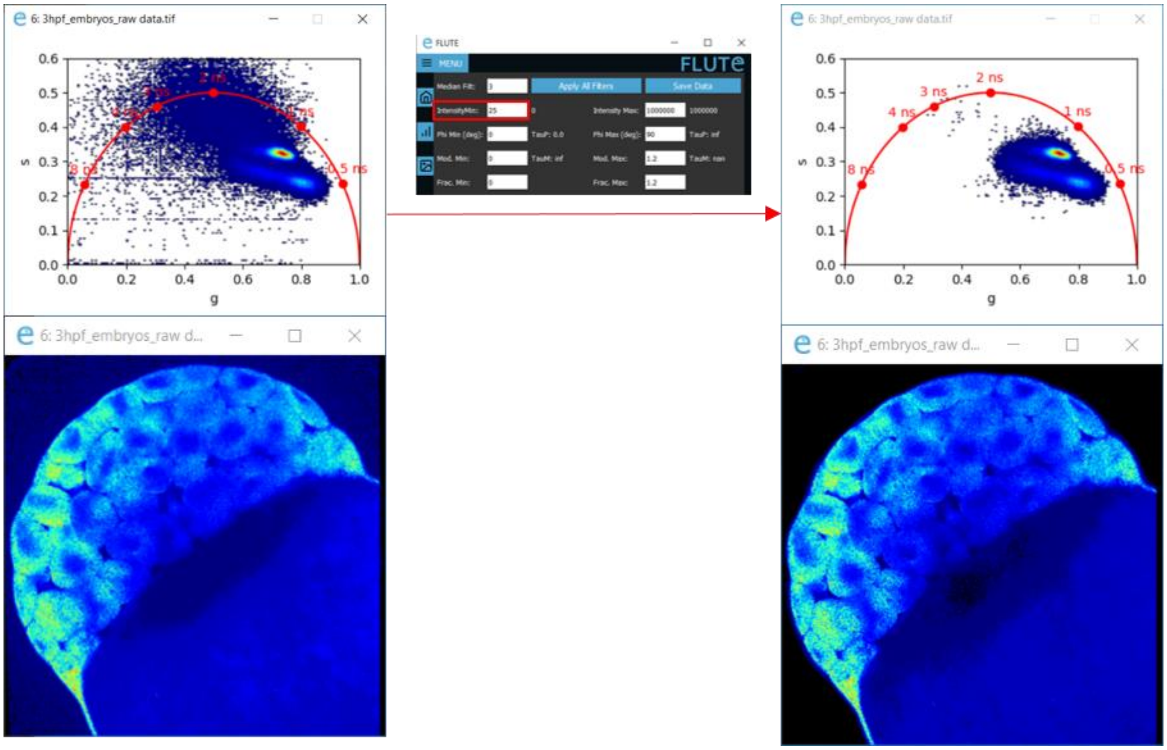


***Figure S8:*** *Applying Intensity threshold to FLIM data.*

### 8. Changing colour maps

FLUTE gives to the user the flexibility to choose the appropriate colour mapping to be applied to the FLIM image and the phasor plot simultaneously. For FLIM data exploration and visualization the image window is switchable between the options presented in figure 9.

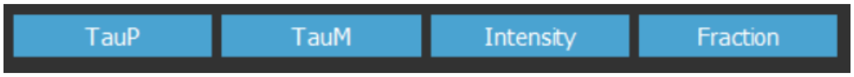


***Figure S9****: Image visualization options: intensity, lifetime contrasts and fraction*

As seen in figure S9, available colour maps are:

- TauP: Phase lifetime calculated with equation 7
- TauM: Modulation lifetime calculated with equation 8
- Intensity: Based on the intensity of the pixel across the entire tiff stack. Both a jet and a greyscale colour maps are available for intensity. Clicking the **Intensity** button will switch between the two.
- Fraction of a molecular species.

The threshold of these colour maps can be adjusted either via the **Interactivity window**, according to figure S10:

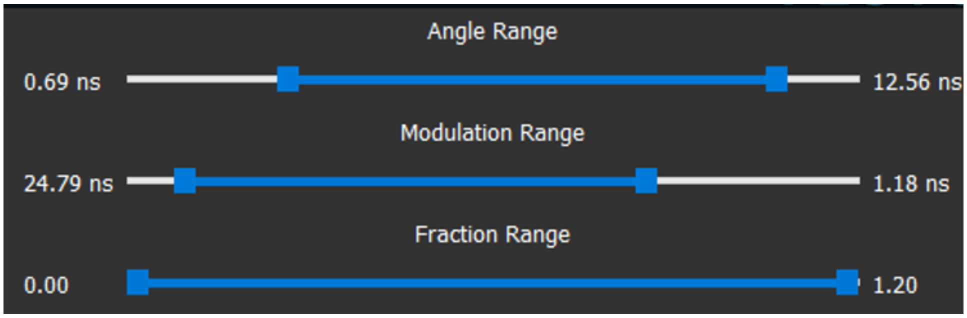


***Figure S10:*** *Thresholding range sliders available in the Interactivity window.*

Or, via the **thresholding window**, shown in figure S11:

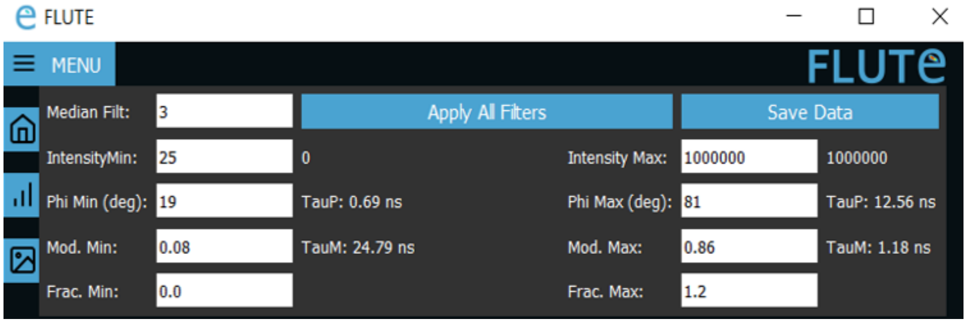


***Figure S11:*** *Thresholding entry boxes available in the Thresholding window.*

#### 8.1 TauP colour map

TauPhase (TauP) is calculated with equation 8. Applying the TauP colour map to the data and setting the **Phi Min (deg)** and **Phi Max (deg)** values to 12 and 35 degrees in the **Thresholding** window respectively will result in the image presented in figure S12.

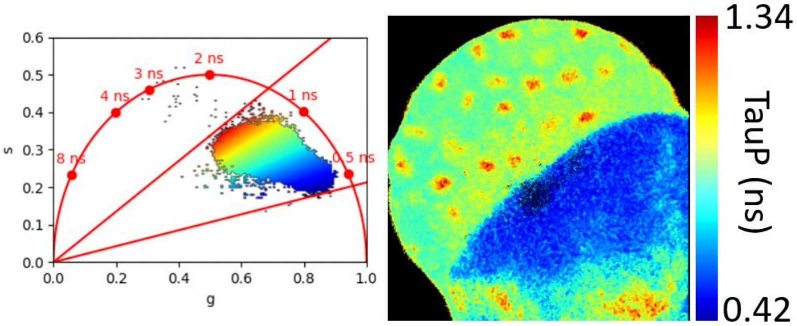


***Figure S12:*** *FLIM data mapped with TauP contrast.*

#### 8.2 TauM colour map

TauModulation (TauM) is calculated with equation 9. Applying the TauM colour map to the data and setting the **Modulation Min** and **Modulation Max** values to 0.59 and 0.92 in the **Thresholding** window respectively will produce the image presented in figure S13.

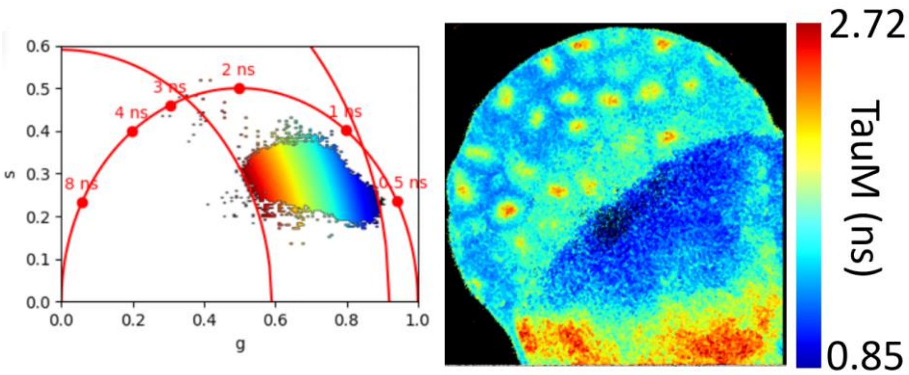


***Figure S13:*** *FLIM data mapped with TauM contrast.*

#### 8.3 Fraction colour map

FLUTE can be used to calculate and map the faction of the molecular species A by inserting the known phasor coordinates of species B. When the fraction colour map is selected, a popup window (figure S14) appears with two options.

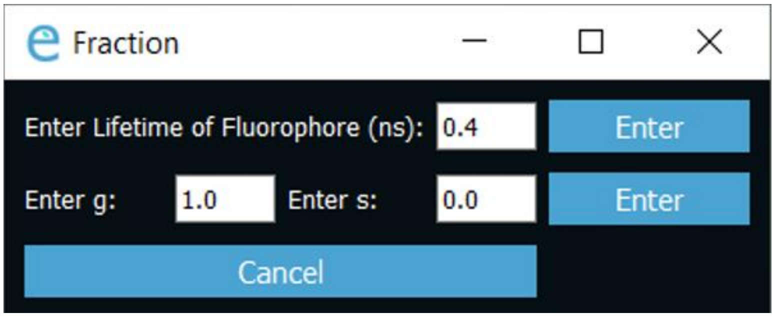


***Figure S14:*** *Entry box to apply options for fraction colour mapping*

**Enter Lifetime of Fluorophore (ns)** will take in account the phasor coordinates of the molecular species B either inserting the single exponential lifetime or the phasor coordinates g and s in the case of a multi-exponential decay. FLUTE then calculates the fraction of the molecular species A with equation 13. For example, the fraction of bound/free NADH can be calculated graphically and mapped in every pixel from the location of the free NADH that has a known single lifetime of 0.4 ns. Figure S15 displays the application of fraction of the bound NADH colour map in the *3hpf_embryos_raw data.tiff* file.

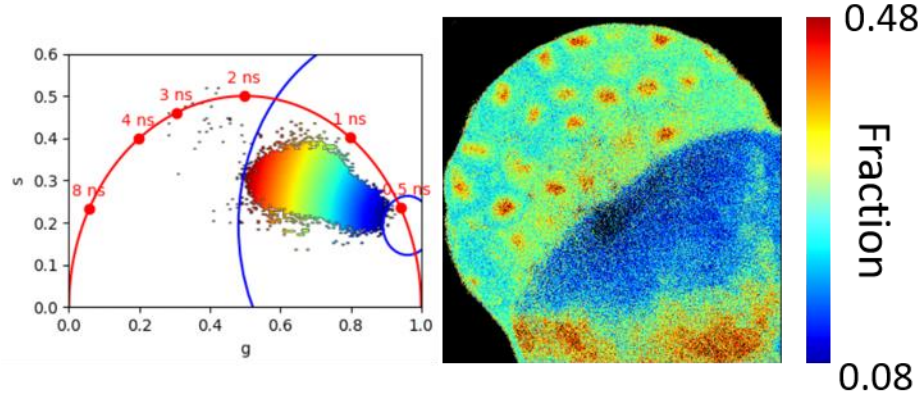


***Figure S15:*** *FLIM Data mapped with fraction of bound NADH contrast.*

#### 8.4 Circle selection

Interactive exploration of the FLIM data can be performed by using multiple coloured cursors of variable sizes to select pixels with similar decays and to highlight simultaneously the corresponding pixels in the image. Areas on the phasor plot can be selected by clicking on an active plot. The modification of colour and size is done through the **Interactivity** window using the **Colour** and **Radius** options. In figure S16, the *3hpf_embryos_raw data.tiff* image is highlighted with a red circle of radius 0.05, and a green circle of radius 0.1, with a greyscale colour map.

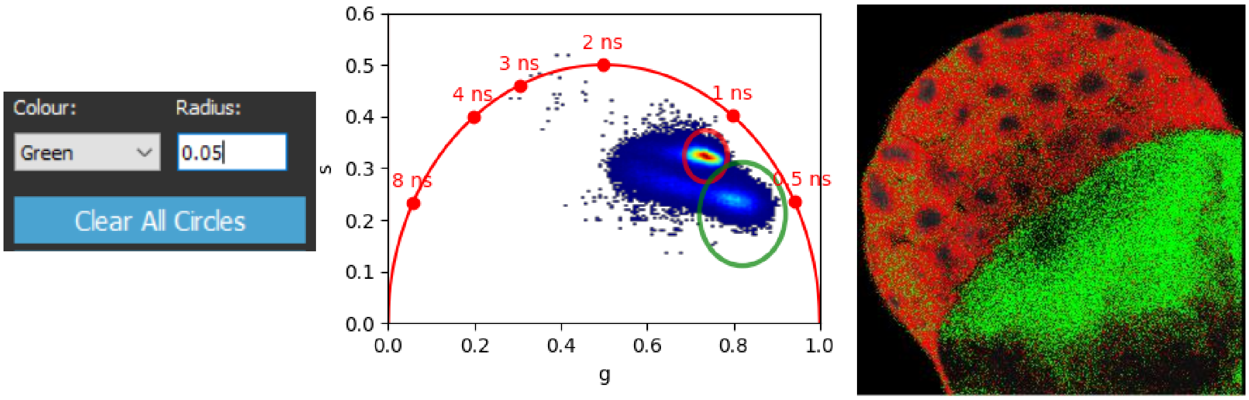


***Figure S16:*** *Data with intensity colour mapping and regions selected with a red circle of radius 0.05 and a green circle of radius 0.1.*

Clicking **Clear All Circles** removes all circles from the plot.

If there are multiple active datasets, clicking on one active phasor plot will draw the same circle on all other active phasor plots.

### 9. Interactivity window

The **Interactivity** window is used for quick and coarse thresholding of the FLIM data and interactive data exploration.

#### 9.1 Range adjustment

Through the interactivity window we can apply a threshold to the phase ($\varphi$), modulation ($m$ ), and fraction range simultaneously using the sliders as displayed in figure S17. For example, the phasor of the *3hpf_embryos_raw data.tiff* image in (figure S17B) has been applied different parameter thresholds that are represented by the coloured lines (red for phase and modulation and blue for fraction). The thresholded pixels are represented in black in the phasor plot.

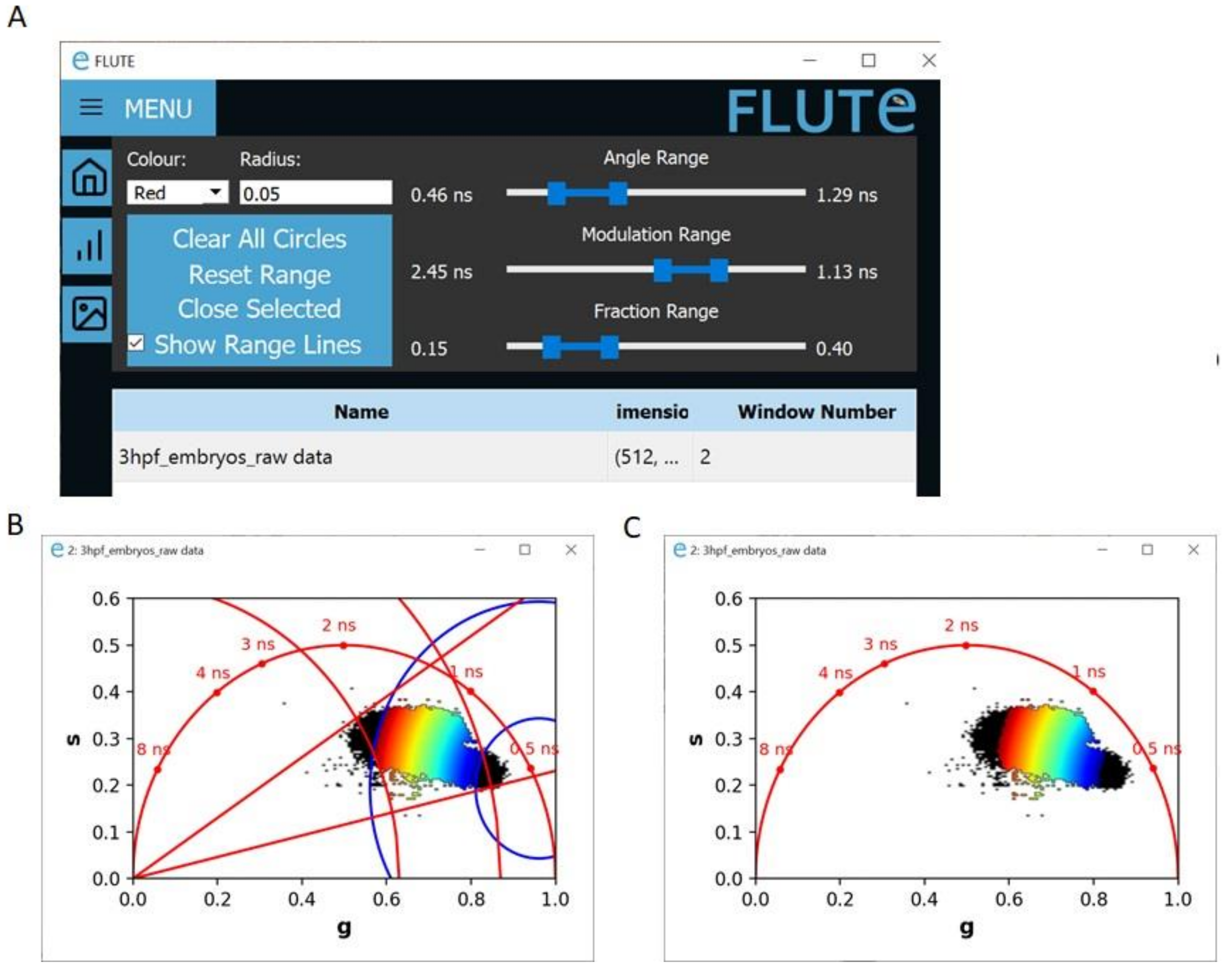


*__Figure S17:__ Range adjustment thought the interactivity window (A) Overview of all available thresholds applied to the data. (B-C) Phasor plots with applied thresholds on the different parameters (phase and modulation and fraction with (B) and without (C) coloured lines that represent the thresholds.*

Clicking **Reset Range** function (Fig.17A) will remove all the applied threshold modifications and will reset the range of the parameters to the default values.

Unclicking **Show Range Lines** function (Fig.17A) will remove all the threshold lines from the phasor plot, while keeping the applied threshold to the FLIM data and phasor cloud (Fig.17C).

#### 9.2 Closing datasets

Closing multiple active datasets is possible by choosing all in the table and clicking **Close Selected**, as shown in figure S18.

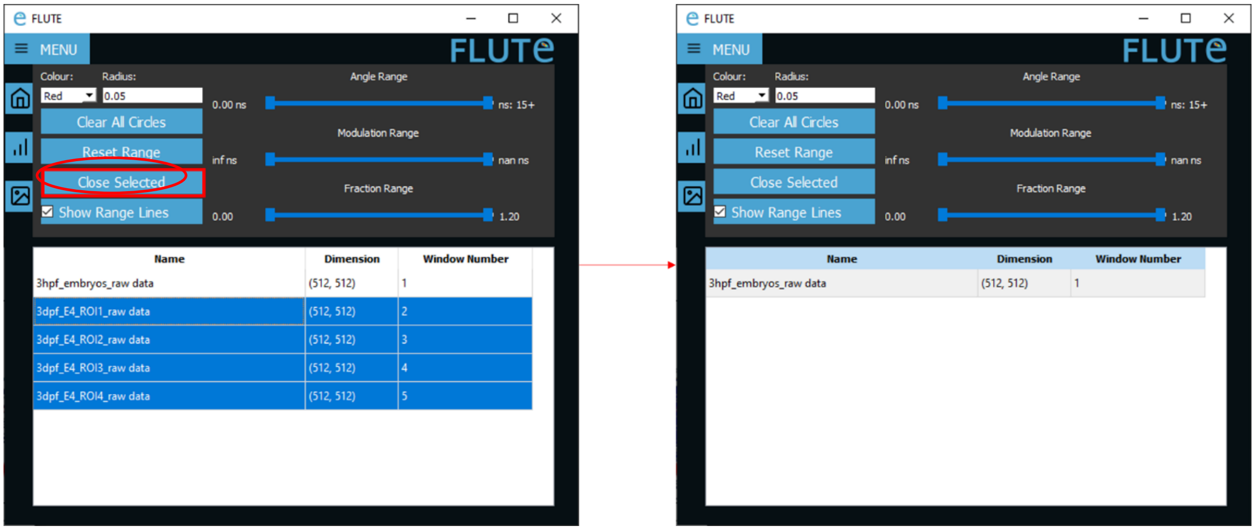


***Figure S18:*** *Closing multiple sets of data at once.*

#### 9.3 Saving data

In addition to fast and interactive FLIM data visualization, analyzed data with applied parameter thresholds can be saved in FLUTE using the Save Data button. Clicking Save Data is followed by a popup window that can save either all the colour maps (All Data) or the colour map that is on display (Currently Displayed). Saved data is listed below; also, an example is given in figure S19.

- …_g.tiff, …_s.tiff: the g and s coordinates with applied parameter thresholds. Thresholded pixels are saved as 'nan'. Data are saved as a 32bit floating point tiff files.
- …_TauP.tiff, …_TauM.tiff, …_Frac.tiff: The TauP lifetime, TauM lifetime, and Fraction values of the dataset with the applied parameter thresholds. Data is saved as a 32bit floating point tiff file with thresholded values saved as 'nan'.
- …_graph_density.png, …_graph _Fraction.png, …_graph _TauM.png, …_graph _TauP.png: Phasor plot screenshots for the various colour maps.
- …_imageFraction.tif, …_imageIntensity.tif, …_imageJet.tif, …_imageTauM.tif, …_imageTauP.tif: screenshots of the FLIM imageswith the respective colour maps with parameter thresholds applied . Data are saved as RGB.
- …_Parameters.txt: List of parameters thresholds applied to the dataset and average values of the phasor coordinates g and s, tauP, tauM and Fraction measured in the mask.

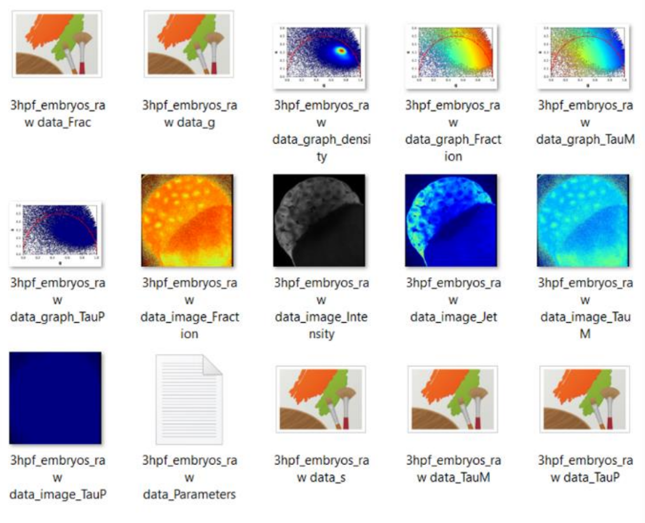


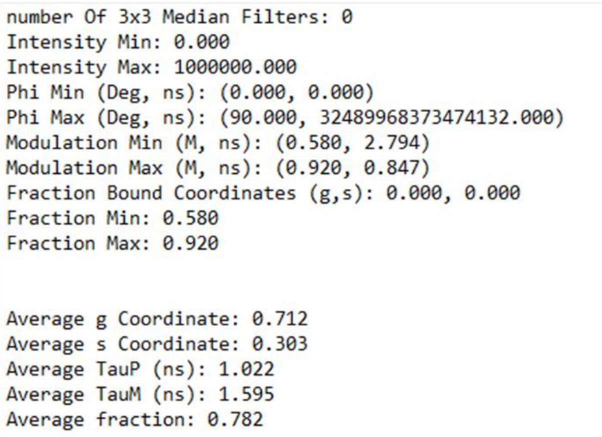
```
number Of 3x3 Median Filters: 0
Intensity Min: 0.000
Intensity Max: 1000000.000
Phi Min (Deg, ns): (0.000, 0.000)
Phi Max (Deg, ns): (90.000, 32489968373474132.000)
Modulation Min (M, ns): (0.580, 2.794)
Modulation Max (M, ns): (0.920, 0.847)
Fraction Bound Coordinates (g,s): 0.000, 0.000
Fraction Min: 0.580
Fraction Max: 0.920

Average g Coordinate: 0.712
Average s Coordinate: 0.303
Average TauP (ns): 1.022
Average TauM (ns): 1.595
Average fraction: 0.782
```

***Figure S19:*** *Example of saved files (left) and parameter file (right)*

### 10. Batch processing

FLUTE has been designed for bulk processing, which is useful for the fast analysis of multiple FLIM data from a single experimental session.

To perform bulk processing, first apply the desired thresholding values in the **Thresholding** window. On the **Home** window, click **Bulk Process**. In the first popup window (figure S20), select the desired data to be processed using ctrl+click or shift+click, and click Open. In the next window, select a folder to save the data. FLUTE will open all datasets; perform the phasor transformation, apply the indicated filters and thresholds, and save all the output data.

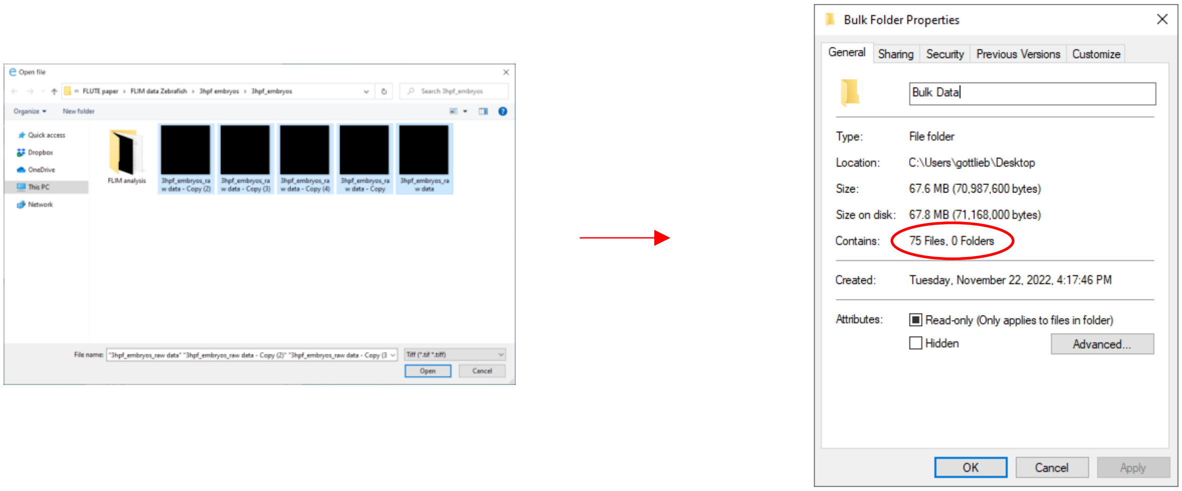


***Figure S20:*** *Batch processing five images leads to 75 saved data files inside the selected folder*
